# Different subregions of monkey lateral prefrontal cortex respond to abstract sequences and their components

**DOI:** 10.1101/2024.02.13.580192

**Authors:** Nadira Yusif Rodriguez, Aarit Ahuja, Debaleena Basu, Theresa H. McKim, Theresa M. Desrochers

## Abstract

Sequential information permeates daily activities, such as when watching for the correct series of buildings to determine when to get off the bus or train. These sequences include periodicity (the spacing of the buildings), the identity of the stimuli (the kind of house), and higher-order more abstract rules that may not depend on the exact stimulus (e.g. house, house, house, business).

Previously, we found that the posterior fundus of area 46 in the monkey lateral prefrontal cortex (LPFC) responds to rule changes in such abstract visual sequences. However, it is unknown if this region responds to other components of the sequence, i.e., image periodicity and identity, in isolation. Further, it is unknown if this region dissociates from other, more ventral LPFC subregions that have been associated with sequences and their components. To address these questions, we used awake functional magnetic resonance imaging in three male macaque monkeys during two no-report visual tasks. One task contained abstract visual sequences, and the other contained no visual sequences but maintained the same image periodicity and identities.

We found the fundus of area 46 responded only to abstract sequence rule violations. In contrast, the ventral bank of area 46 responded to changes in image periodicity and identity, but not changes in the abstract sequence. These results suggest a functional specialization within anatomical substructures of LPFC to signal different kinds of stimulus regularities. This specialization may provide key scaffolding to identify abstract patterns and construct complex models of the world for daily living.

**Significance Statement:** Daily tasks, such as a bus commute, require tracking or monitoring your place (same, same, same, different building) until your stop. Sequence components such as rule, periodicity (timing), and item identity are involved in this process. While prior work located responses to sequence rule changes to area 46 of monkey lateral prefrontal cortex (LPFC) using awake monkey fMRI, less was known about other components. We found that LPFC subregions differentiated between sequence components. Area 46 posterior fundus responded to abstract visual sequence rule changes, but not to changes in image periodicity or identity. The converse was true for the more ventral, adjacent shoulder region. These results suggest that interactions between adjacent LPFC subregions provide key scaffolding for complex daily behaviors.

## Introduction

When commuting to work, such as when taking a bus, you may internally track house, house, house, store (same, same, same, different building) until you arrive at your stop. This recognition illustrates an essential process: the monitoring of abstract sequences. These sequences are abstract because they do not depend on the identity of the individual stimuli (e.g., changing the color of the house). The same system that facilitates sequential tracking also enables the detection of changes or deviations to an existing sequence. Sequences are defined by multiple elements that we specifically refer to as components. These sequential components include item identity, periodicity (temporal structure), and rule. Abstract sequence rule deviations may encompass changes in these individual sequence components. How does the brain track changes to sequences and their components?

We identified a brain region that responds to abstract sequence deviations, but whether the same or other regions also monitor changes in sequential components has not been tested. Previously, we identified a specific subregion within the fundus of posterior area 46 of the lateral prefrontal cortex (LPFC) as uniquely responding to sequential changes using awake monkey functional magnetic resonance imaging (fMRI) (Yusif Rodriguez et al., 2023). Beyond the rule that these abstract sequences followed, they had two (previously mentioned) main components: image identity and periodicity. These components were controlled for when determining responses to infrequent changes in the abstract visual sequence. However, LPFC is also known to respond to changes in image identity or periodicity. Responses to infrequent (sometimes referred to as “oddball”) stimuli have been reported in monkey LPFC using an array of techniques (Chao et al., 2018; Camalier et al., 2019; Grohn et al., 2020). Changes in stimulus periodicity could also elicit responses in LPFC because responses in LPFC are modulated by the duration preceding the auditory or visual stimulus (Onoe et al., 2001; Genovesio et al., 2006; Chiba et al., 2021). Therefore, the question arises as to whether the same, abstract sequence coding, or other LPFC subregions respond to changes in image periodicity or identity alone.

The ventral LPFC (VLPFC) is a prime candidate as another subregion within the LPFC that could respond to sequential components. FMRI studies in macaques have shown VLPFC activity during auditory sequential tasks and sequence deviants (Wang et al., 2015; Vergnieux and Vogels, 2020). Studies using electrophysiology provided evidence for the representation of generalizable sequential structures and changes to these structures in neuronal population responses within VLPFC (Esmailpour et al., 2023; Bellet et al., 2024). VLPFC also responds to non-sequential information that shares similar features with sequential tasks, such as prediction error (Uhrig et al., 2014; Chao et al., 2018) and responses to infrequent (“oddball”) items (Uhrig et al., 2014; Suda et al., 2022). VLPFC is often observed as more directly representing sensory visual information (compared to DLPFC, which can be more spatial or action oriented).

Responses in VLPFC for non-spatial object-based features include color, shape, and object type (Meyer et al., 2011; Yamagata et al., 2012; Tang et al., 2021; Xu et al., 2022). Thus, there is evidence to suggest that VLPFC could respond to sequential components and abstract sequences, underscoring the importance for dissociating between LPFC subregions.

To dissociate between LPFC subregions, we defined a more ventral, yet adjacent brain area. Historically, many conventions have been applied to naming subregions within the LPFC (Walker, 1940; Petrides and Pandya, 1999; Rapan et al., 2023). While some studies refer to VLPFC as the region ventral to the arcuate sulcus, primarily Brodmann area 44 and potentially parts of 6VR (Rapan et al., 2023), others refer to the region ventral to the principal sulcus and dorsal to the arcuate sulcus that can include Brodmann areas 46, 9/46, 45A, and 45B along with more anterior territory such as Brodmann areas 47 and 12 (Rapan et al., 2023). Our purpose here is not to adjudicate among the definitions, but take advantage of a commonality that, in general, more ventral regions of the LPFC are more biased towards sensory, object-based, or non-spatial information (Meyer et al., 2011; Yamagata et al., 2012; Tang et al., 2021; Xu et al., 2022). Therefore, an area that is more ventral, yet adjacent to the subregion in area 46 where we previously observed sequential responses is an ideal candidate for comparison. A region defined as thus would not make assumptions about naming conventions and still be clearly within the LPFC. Using this region, we determined if there were differences in abstract sequence or component representation between the two LPFC sub-regions.

To address the questions of whether LPFC subregions dissociate based on 1) responses to abstract sequences, and 2) responses to sequence components (image identity and periodicity), we conducted an awake monkey fMRI experiment. Monkeys performed two no-report tasks, one that contained abstract visual sequences (previously reported on in Yusif Rodriguez et al. (2023)), and another that did not contain abstract visual sequences but maintained the image identities and timing structure of the sequence task. We defined two LPFC subregions using a parcellation of the PFC (Rapan et al., 2023): the posterior fundus of area 46 (p46f) that overlapped with the previously identified sequence responsive subregion (Yusif Rodriguez et al., 2023), and the adjacent posterior ventral shoulder of area 46 (p46v). Building on our previous observations, we hypothesized that p46f would show responses unique to changes in abstract visual sequences and not to their image identity and periodicity components. We hypothesized the contrary for p46v: responses to changes in sequence components but not sequences themselves. Our results broadly supported these hypotheses, with sequence responses in p46f and not p46v, and responses to image identity and periodicity in p46v. These results further our understanding of the representation of abstract visual sequences in adjacent subregions in the LPFC.

## Materials and Methods

### Participants

We tested three adult male rhesus macaques (ages spanning 6-12 years during data collection, 9-14 kg). All procedures followed the NIH Guide for Care and Use of Laboratory Animals and were approved by the Institutional Animal Care and Use Committee (IACUC) at Brown University.

### Task Design and Procedure

All visual stimuli used in this study were displayed using an OpenGL-based software system developed by Dr. David Sheinberg at Brown University. The experimental task was controlled by a QNX real-time operating system using a state machine. Eye position was monitored using video eye tracking (Eyelink 1000, SR Research). Stimuli were displayed at the scanner on a 24-inch BOLDscreen flat-panel display (Cambridge Systems).

Each image presentation consisted of fractal stimulus (approximately 8° visual angle) with varying colors and features. Fractals were generated using MATLAB for each scanning session using custom scripts based on stimuli from Kim and Hikosaka (2013) following the instructions outlined in Miyashita et al. (1991). For each scan session, new, luminance matched fractal sets were generated. All stimuli were presented on a gray background, with a fixation spot that was always present on the screen superimposed on the images. To provide behavioral feedback, the fixation spot was yellow when the monkey was successfully maintaining fixation and red if the monkey was not fixating. Stimuli were displayed for 0.1, 0.2, or 0.3 s each, depending on the task, sequence type, and timing template.

The timing of liquid rewards was the same across tasks and not contingent on image presentations, only on the monkey maintaining fixation. Rewards were delivered on a graduated schedule such that the longer the monkey maintained fixation, the more frequent rewards were administered (Leite et al., 2002). The first reward was given after 4 s of continuous fixation. After two consecutive rewards of the same fixation duration, the fixation duration required to obtain reward was decreased by 0.5 s. The minimum duration between rewards that the monkey could obtain was 0.5 s. Fixation had to be maintained within a small window (typically 3° of visual angle) around the fixation spot to not break fixation. The only exception was a brief time window (0.32 s) provided for blinks. If the monkey’s eyes left the fixation window and returned within that time window, it would not trigger a fixation break. If fixation was broken, the reward schedule would restart at the maximum 4 s duration required to obtain reward.

Tasks were organized into runs. Runs typically lasted approximately 10 min and only one task was shown for each run. The order of tasks was pseudo-randomized within a scanning session (one day) to balance the overall number of runs for each task and their presentation order. Monkeys completed approximately 10 runs in a session.

Runs were initiated according to the monkey’s fixation behavior to ensure that the monkey was not moving and engaged in the task before acquiring functional images. During this pre-scan period, a fixation spot was presented. Once the monkey successfully acquired this fixation spot and received approximately four liquid rewards (12 – 16 s), functional image acquisition and the first habituation block were initiated. Monkeys maintained fixation for the duration of the run.

### Abstract Sequence Viewing (SEQ) Task

The details of the abstract sequence viewing task have been previously described (Yusif Rodriguez et al., 2023) and are briefly summarized here. There were a total of five sequence types and nine timing templates (**Figure 1**). The inter-sequence interval was jittered to decorrelate across timing templates (mean 2 s, 0.25-8 s).

**Figure 1.**
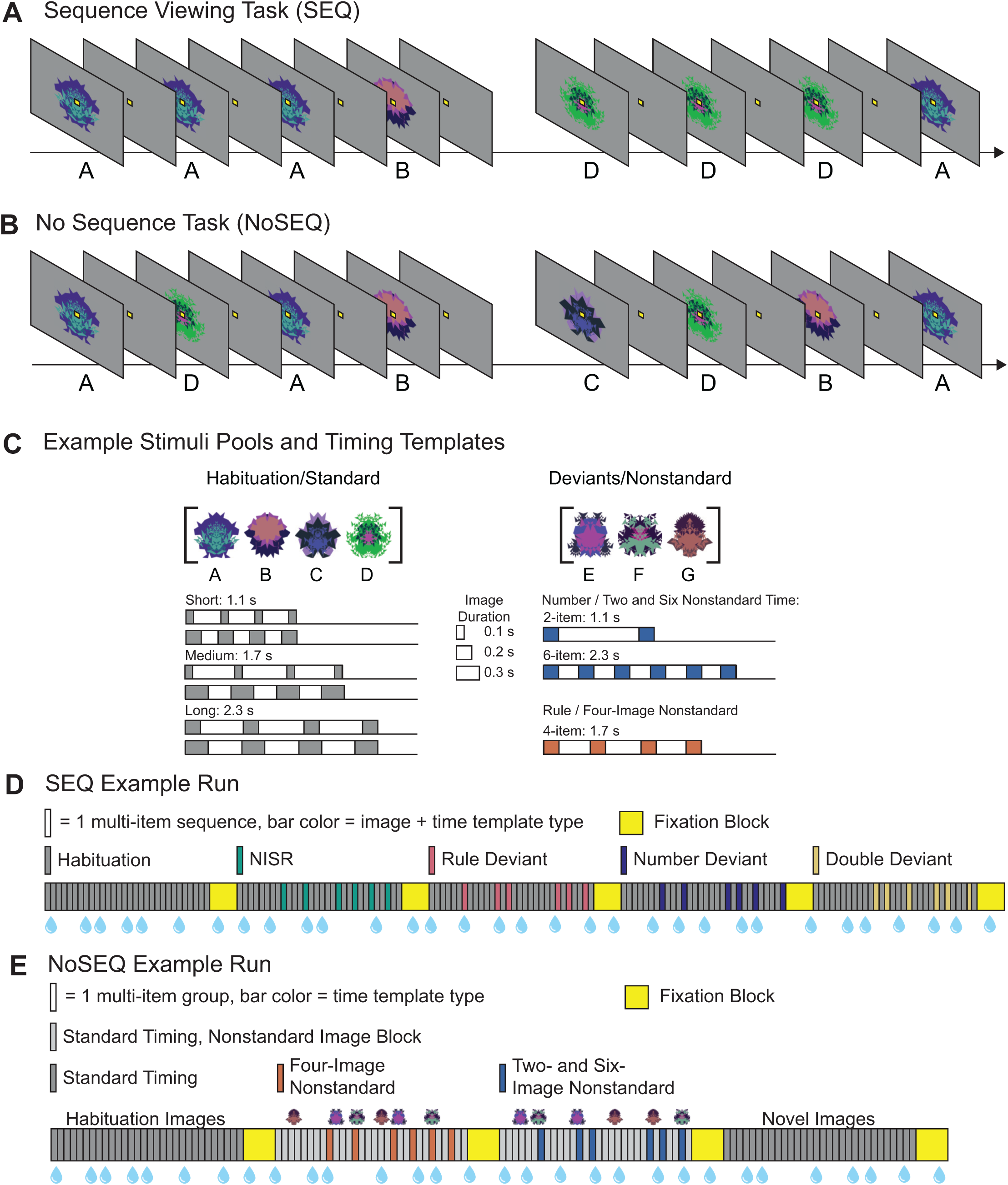
Sequence viewing task (SEQ) and No Sequence (NoSEQ) task structure. Both tasks are no-report. The monkey maintains fixation at the central fixation spot throughout both tasks. **A**. Example partial habituation block from SEQ task for sequence rule three same, one different (AAAB) and habituation timing templates. **B**. Example partial standard block from NoSEQ illustrating non-sequential image order and standard timing templates. **C**. Example stimuli pools (top) show a set of images that would be used in a single scanning session for both tasks (but termed differently depending on the task). NoSEQ additionally contains a novel images category, with different images not exemplified here. New images are used each session. Six possible habituation/standard event timing templates (bottom, left) and deviant/nonstandard event timing templates (bottom, right) illustrated with gray rectangles indicating individual image presentations. Total sequence/grouping durations are listed for each template type. **D**. Example SEQ run, with each bar indicating one multi-image sequence: four images in habituation, new items same rule (NISR), and rule deviants; two or six images in number and double deviants. Each block contains 30 sequences. The first block contains only habituation sequences and subsequent blocks (order counterbalanced) contain only one of the four deviant types in six out of the 30 sequences. **E.** Example NoSEQ run where there is no sequential order to the displayed images. Each bar indicates a multi-image set grouped by timing template and each block contains 30 image groupings. To parallel SEQ structure, the first block in NoSEQ contains only standard images and timing templates. In the two following blocks (order counterbalanced), six out of the 30 groupings have nonstandard timing templates. One block has 4-image nonstandard timing (4NST) that parallels the rule deviant block in SEQ, and the other block has 2– and 6-image nonstandard timing (2/6NST) together to parallel the number deviant block in SEQ. Also, in the nonstandard blocks, 20% of the fractal images shown (in pseudorandom order) are nonstandard (indicated by miniaturized nonstandard fractals), and the rest standard, again to mirror the proportions in SEQ. Task relevant blocks alternate with fixation blocks for both SEQ and NoSEQ tasks. In fixation blocks, monkeys maintained fixation on the fixation spot while no other images were displayed. Blue water droplets schematize reward delivery, which is decoupled from sequence viewing and delivered on a graduated schedule based on the duration the monkey has maintained fixation.

#### Habituation Sequences

Habituation sequences used images drawn from a pool of four fractals [A, B, C, D] and were arranged to follow one of two possible rules: three the same, one different, and four the same. All four-image sequences used one of six possible timing templates (**Figure 1C**).

#### Deviant Sequences

Deviant sequences used images drawn from a different pool of three fractals [E, F, G]. All deviant images were displayed for 0.2 s and used the same general timing template (adjusted for the number of items in the sequence). There were four deviant types, as follows:

● *New Items, Same Rule (NISR)*: four-image sequences that follow the same rule as the habituation sequences.
● *Rule Deviants*: four-image sequences that follow the alternate rule not used in the habituation sequences.
● *Number Deviants*: two-or six-image sequences that follow the same rule as the habituation sequences.
● *Double Deviants*: combine the rule and number deviant types and contain two-or six-image sequences that follow the alternate rule not used in the habituation sequences.

#### Block Structure

All blocks contained 30 sequences and an equal number of the six possible timing templates for habituation sequences. Sequences could not start with the same image as the final fractal of the previous sequence. In deviant blocks, six of the 30 sequences were deviant sequences. Deviant sequences did not occur in the first six sequences (to avoid block initiation) or consecutively. Blocks with two– and six-image sequences contained an equal number of both.

#### Run Structure

Each run contained five sequence blocks interleaved with 14 s fixation blocks, during which only a fixation spot was present with no additional visual stimuli (**Figure 1D**). Monkeys maintained fixation throughout the sequence and fixation blocks. The first sequence block was always all habituation sequences. The four subsequent sequence blocks each contained one type of deviant sequence. The sequential rule used for each run was counterbalanced across runs and sessions to have an equal number of each. Monkeys typically completed 4-8 runs of this task (among other tasks) in a session.

### No Sequence (NoSEQ) Task

The main difference between NoSEQ and SEQ was that images were not arranged according to a sequential rule (as in SEQ) and instead displayed in pseudorandom order such that there were no consecutively repeated images. Images were still displayed as grouped into four-, two-, or six-image sets, depending on the block. All the remaining basic structure of NoSEQ was the same as SEQ. We adjusted the terminology to reflect this fact and more clearly dissociate between the tasks. There were the same 9 timing templates in NoSEQ as in SEQ. However, for NoSEQ, rather than referring to them as 6 habituation and 3 deviant timing templates as in SEQ, we refer to them as 6 standard and 3 nonstandard timing templates in NoSEQ. Similarly, the same two pools of fractal images referred to as habituation and deviant in SEQ are referred to as the 4 standard and 3 nonstandard images in NoSEQ. We underscore that the timing templates and image pools are the same between SEQ and NoSEQ within a single scanning session, despite the difference in naming.

#### Block Types and Structure

We define trials as a series of grouped images with the same timing structure. Most trials contained four images, with some containing two or six (described further below). All blocks contained 30 trials (120 images total). We note that even though images are grouped into trials, reward is on an independent schedule based on the duration of fixation, as in the SEQ task. The first six trials in a block did not contain nonstandard timing templates or nonstandard images.

Each block contained key differences with respect to the composition of the timing templates and images used. There were four possible block types (**Figure 1E**), as follows:

● *All Standard Timing*: Each four-image trial used one of the 6 possible standard timing templates (5 of each). Images were drawn only from the standard pool. This condition is the same structure as habituation timing in the SEQ task.
● *Four-Image Nonstandard Timing (4NST)*: Six trials had four-image nonstandard timing and the remaining 24 trials had standard timing. This timing structure matched the NISR and rule deviant blocks in the SEQ task. The relative fraction of nonstandard images matched the SEQ task (20%, 24 individual images), but they were randomly intermixed with images from the standard pool.
● *Two– and Six-Image Nonstandard Timing (2/6NST)*: Six trials had two-or six-image nonstandard timing (three of each) and the remaining 24 trials had standard timing. This timing structure matched the number deviant blocks in the SEQ task. As in Four-Image Nonstandard Timing blocks, 20% of images were drawn from the nonstandard image pool and the remainder from the standard image pool. All images were displayed in random order.
● *Novel*: As in the Standard Timing block, each four-image trial used one of the 6 possible standard timing templates (5 of each). However, the images came from a novel pool of four images that had not been used in either the standard or nonstandard image pools.

#### Run Structure

Each run was composed of four image blocks, interleaved with 14 s fixation blocks. As in the SEQ task, fixation blocks consisted of only a fixation spot present and no additional visual stimuli where the monkey had to maintain fixation. The first block of each run was always an All Standard Timing block. The two subsequent blocks were either a Four-Image Nonstandard Timing block or a Two– and Six-Image Nonstandard Timing block, with their order counterbalanced across runs. The last block was always a Novel block. Runs lasted approximately 10 min. Monkeys typically completed 2-4 runs of this task (among other tasks) in a single scanning session.

## Data Acquisition

### FMRI Data Acquisition

Methods are as described in (Yusif Rodriguez et al., 2023) and briefly summarized here. Monkeys sat in the “sphinx” position in an MR-safe primate chair (Applied Prototype, Franklin, MA or custom-made by Brown University) with their head restrained using a plastic “post” (PEEK, Applied Prototype, Franklin, MA) affixed to the monkeys’ head and the primate chair. Monkeys wore earplugs during MRI scanning (Mack’s Soft Moldable Silicone Putty Ear Plugs, “kid’s” size). Monkeys were habituated to all scanning procedures prior to the MRI scanning sessions.

Approximately 30-60 min prior to each scanning session, monkeys were intravenously injected with a contrast agent: monocrystalline iron oxide nanoparticle (MION, Feraheme (ferumoxytol), AMAG Pharmaceuticals, Inc., Waltham, MA, 30 mg per mL or BioPal Molday ION, Biophysics Assay Lab Inc., Worcester, MA, 30 mg per mL). MION was injected into the saphenous vein below the knee (7 mg/kg), then flushed with a volume of sterile saline approximately double the volume of the MION injected. No additional MION was added during scanning.

A Siemens 3T PRISMA MRI system with a custom six-channel surface coil (ScanMed, Omaha, NE) at the Brown University MRI Research Facility was used for whole-brain imaging. Anatomical scans consisted of a T1-MPRAGE (repetition time, TR, 2700 ms; echo time, TE, 3.16 ms; flip angle, 9°; 208 sagittal slices; 0.5 x 0.5 x 0.5 mm), a T2 anatomical (TR, 3200 ms; TE 410 ms; variable flip angle; 192 interleaved transversal slices; 0.4 x 0.4 x 0.4 mm), and an additional high resolution T2 anatomical (TR, 8020 ms; TE 44 ms; flip angle, 122°; 30 interleaved transversal slices; 0.4 x 0.4 x 1.2 mm). Functional images were acquired using a fat-saturated gradient-echoplanar sequence (TR, 1.8 s; TE, 15 ms; flip angle, 80°; 40 interleaved axial slices; 1.1 x 1.1 x 1.1 mm).

## Data Analysis

All preprocessing and data inclusion criteria are the same as in (Yusif Rodriguez et al., 2023). Most analyses were performed in Matlab using SPM 12 (https://www.fil.ion.ucl.ac.uk/spm). Prior to analysis, data were preprocessed using the following steps: reorienting (to ensure proper assignment of the x,y,z planes), motion correction (realignment), normalization, and spatial smoothing (2 mm isotropic Gaussian kernel separately for gray matter and white matter). All steps were performed on individual runs separately. The T1-MPRAGE anatomical image was skull stripped using FSL BET brain extraction tool (http://www.fmrib.ox.ac.uk/fsl/) to facilitate normalization. All images were normalized to the 112-RM SL macaque atlas (McLaren et al., 2009).

Runs were included for analysis only if they met the following criteria: the monkey had to be performing well and a sufficient number of acquisition volumes within the run had to pass data quality checks. The monkey’s performance was evaluated by calculating the percentage of time within a run that fixation was maintained. Runs were excluded if the monkey was fixating < 80% of the time. We used the ART toolbox (Artifact Detection Tools, https://www.nitrc.org/projects/artifact_detect) to detect outlier volumes (standard global mean; global signal detection outlier detection threshold = 4.5; motion threshold = 1.1mm; scan to scan motion and global signal change for outlier detection). Any run with greater than 12% of volumes excluded was excluded from analysis (**Table 1**).

**Table 1.** Data excluded and included for analysis.

| Percent Excluded Fixation |  |  |  |  |
| --- | --- | --- | --- | --- |
|  | Monkey B | Monkey J | Monkey W |  |
| SEQ | 6.89% | 10.46% | 3.03% |  |
| NoSEQ | 6.96% | 10.13% | 5.06% |  |
| Percent Excluded Motion |  |  |  |  |
|  | Monkey B | Monkey J | Monkey W |  |
| SEQ | 15.15% | 0% | 0.6% |  |
| NoSEQ | 16.5% | 0.63% | 1.89% |  |
| Total Included Runs |  |  |  |  |
|  | Monkey B | Monkey J | Monkey W | Total Runs |
| SEQ | 70 | 65 | 97 | 232 |
| NoSEQ | 17 | 38 | 43 | 98 |

### FMRI Models

For all models, data were binned to evenly distribute included runs from the SEQ and NoSEQ tasks (**Table 1**) into pseudo-subject bins. Each bin contained data from only one monkey and distributed runs from the SEQ and NoSEQ tasks as evenly as possible. Each bin contained approximately 20 SEQ and 10 NoSEQ runs. Runs from earlier and later scanning sessions were pseudorandomly distributed across bins. For the SEQ task, both rule types (AAAA and AAAB) were evenly distributed in each bin. This binning procedure resulted in 11 total pseudo-subject bins. Of the 11 pseudo-subject bins, 5 were monkey W, 4 were monkey J, and 2 were monkey B.

Within-subject statistical models were constructed under the assumptions of the general linear model (GLM) in SPM 12 for each pseudo-subject bin. Condition regressors were all convolved with a gamma function (shape parameter = 1.55, scale parameter = 0.022727) to model the MION hemodynamic response function (Vanduffel and Farivar, 2014). The first six image groups (24 images) and reward times were included as nuisance conditions. Additional nuisance regressors were included for the six motion estimate parameters (translation and rotation) and image variability (standard deviation of within-run image movement variability, calculated using the ART toolbox). Outlier volumes determined with the ART toolbox in preprocessing were “scrubbed” by adding an additional regressors, each with a “1” only at the volume to be excluded. The equation for the GLM is below (Poline and Brett, 2012):

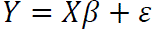

Where *Y* is the (*n*,1) time series data (*n* = number of time points or scans), *X* is (*n*, *p*) design matrix of regressors, *β* is the vector of parameters, and *ε* is the error vector. Regressors included are the 20 listed in **Table 2** plus the nine nuisance regressors listed above (total of 29 regressors) and any additional columns required for “scrubbing” (one regressor per volume scrubbed). The baseline used for comparisons was implicit in that it included unmodeled time for which there were no explicitly defined condition, nuisance, or “scrubbed” regressors (i.e., fixation only time where there were no images displayed during fixation only blocks and inter-sequence intervals).

**Table 2.** Regressors used in SEQ and NoSEQ model. Both tasks are modeled together.

SEQ Task Regressors
| First Block | Subsequent Blocks |
| --- | --- |
| Habituation Short | Habituation Short |
| Habituation Medium | Habituation Medium |
| Habituation Long | Habituation Long |
|  | New Item Same Rule (NISR) |
|  | Rule Deviant |
|  | Number Deviant – 2 items |
|  | Number Deviant – 6 items |
|  | Double Deviant – 2 items |
|  | Double Deviant – 6 items |

NoSEQ Task Regressors
| First Block | Subsequent Blocks |
| --- | --- |
| Standard timing - Short | Standard timing - Short |
| Standard timing - Medium | Standard timing - Medium |
| Standard timing - Long | Standard timing - Long |
|  | Nonstandard image |
|  | 4-image nonstandard timing |
|  | 2-image nonstandard timing |
|  | 6-image nonstandard timing |
|  | Novel image, standard timing - Short |
|  | Novel image, standard timing - Medium |
|  | Novel image, standard timing - Long |

Regressors were estimated using a bin-specific fixed-effects model. Whole-brain estimates of bin-specific effects were entered into second-level analyses that treated bin as a random effect. One-sample t-tests (contrast value vs zero, p < 0.005) were used to assess significance. These effects were corrected for multiple comparisons when examining whole-brain group voxelwise effects using extent thresholds at the cluster level to yield false discovery rate (FDR) error correction (p < 0.05).

To assess the univariate effects of deviant sequences, we constructed a general linear model (GLM) using instantaneous stimulus onset regressors. Both tasks were modeled simultaneously, with runs from both tasks included in each pseudo-subject bin. For the SEQ task, onsets were modeled similarly as described in Yusif Rodriguez et al. (2023). Onsets were modeled at the first item in each sequence type. Habituation and deviant sequences were modeled separately. Habituation sequences were divided by timing template (short, medium, and long) and whether they came from the first block containing only habituation sequences or a subsequent block that contained deviant and habituation images, yielding six total habituation sequence regressors. Deviant sequences were modeled separately according to their type: NISR, rule deviants, number deviants (two– and six-image), and double deviants (two– and six-image), yielding six total deviant sequence regressors. In total, the SEQ task contained 12 condition regressors (**Table 2**).

For the NoSEQ task, onsets were modeled for the first item in each group of images (a single timing template). Standard and nonstandard timing templates were modeled separately. As in the SEQ task, standard timing templates were divided by those occurring in the first block (where there were no nonstandard timing templates or images) and those occurring in subsequent blocks that contained nonstandard images and timing templates. Standard timing templates were again divided by short, medium, and long yielding a total of six standard timing template regressors. Nonstandard timing templates were modeled separately as four-image and two– and six-image, yielding three total nonstandard timing template regressors. Nonstandard images that were randomly interspersed in blocks that contained nonstandard timing templates were modeled separately at the onset of each individual nonstandard image. The novel image block was also separately modeled and divided by the three standard timing templates (short, medium, long); however, these were not included in analyses. In summary, the NoSEQ task contained six standard time, three nonstandard time, one nonstandard image, and three novel image regressors for a total of 13 regressors (**Table 2**).

### ROI Analysis

Individual area 46 subregion ROI images were directly acquired from the MEBRAINS Multilevel Macaque Atlas (Balan et al., 2024) (https://www.ebrains.eu/tools/monkey-brain-atlas). Individual subregion image warps were created from their native space to 112RM-SL space using Rhemap (Sirmpilatze and Klink, 2020) (https://github.com/PRIME-RE/RheMAP). Individual warps were then applied to create images used in ROI analysis for p46v (**Figure 2**). Because the ROI used in Yusif Rodriguez et al. (2023) spanned subregions p46df and p46vf and responses in these subregions were not distinct, for simplicity, we combined subregions in the fundus of area 46 to create p46f (p46df + p46vf).

**Figure 2.**
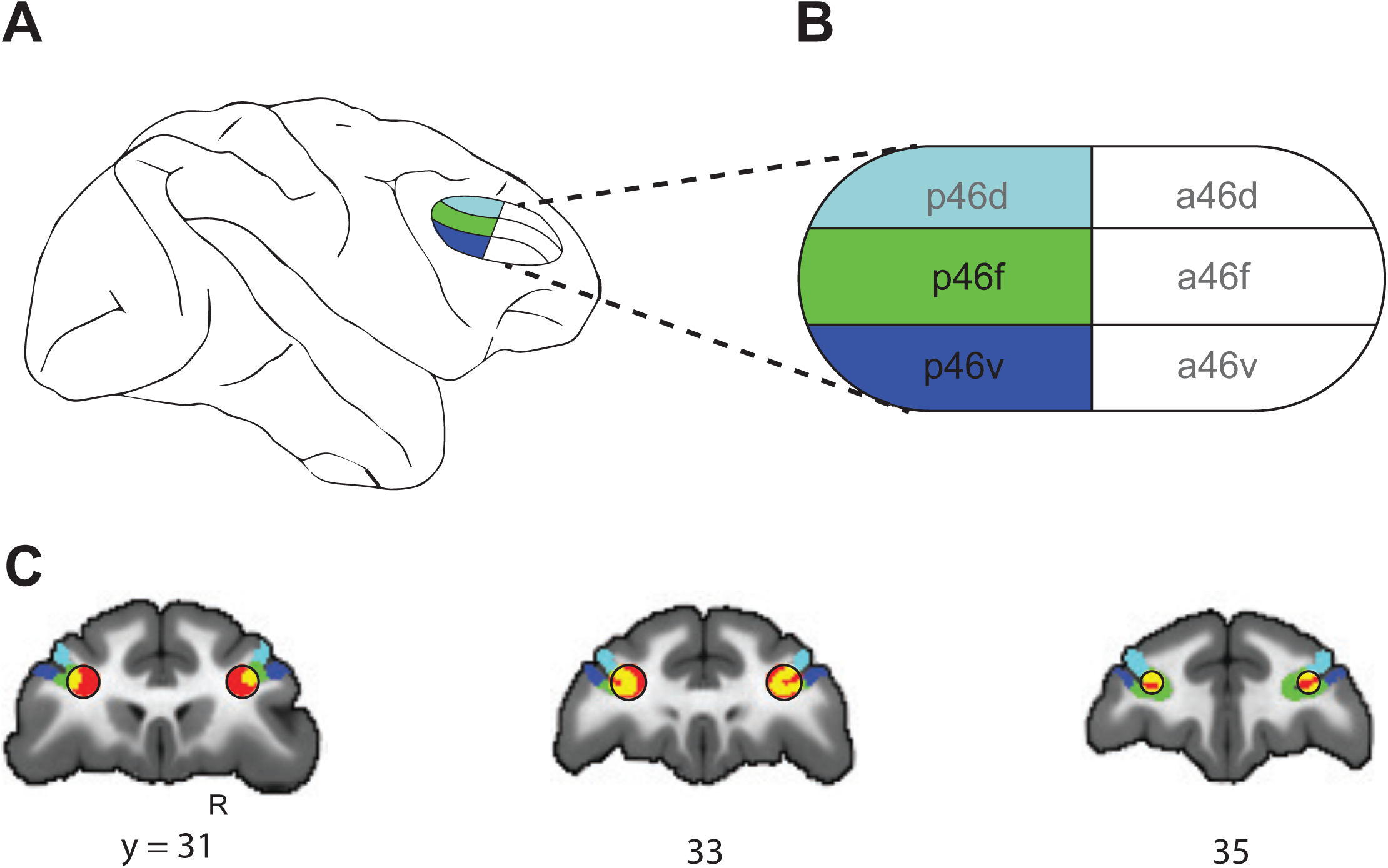
Schematic of anatomical subdivisions of area 46. **A.** Schematic of the area 46 subregions in the LPFC based on the atlas by Rapan et al. (2023) depicted on the right lateral surface of the macaque brain. **B.** Same cortical subregions illustrated in **A** with areas of comparison p46f and p46v in green and blue. Area p46d (light blue) shown only for context, indicated by gray lettering. **C.** Coronal slices displaying the area 46 ROI sphere used in Yusif Rodriguez et al. 2023 (red, outlined in black) superimposed on area 46 subregions. ROIs used for analysis in this study were green and blue, corresponding to regions illustrated in **A**). Light blue p46d only included for comparison and not used in analyses. Yellow voxels indicate overlap between the previous red sphere and current p46f ROI (green).

To compare activation within and across ROIs in a manner that controlled for variance, we extracted average t-values for each ROI from whole-brain t-maps using the Marsbar toolbox (Jean-Baptiste Poline, 2002). T-maps were the result of voxel-wise t-tests of parameter weights for the conditions of interest scaled by the residual error in the model (GLM) compared to a null hypothesis of zero, i.e., the resulting t-values from the condition > baseline contrast (see Poline & Brett (2012) for further details). This procedure resulted in a total of 11 t-values for each condition (one for each pseudo-subject bin: n = 11 bins) that were entered into repeated measures analyses of variance (RM-ANOVAs) with the identity of the monkey entered as a covariate.

## Results

We show awake fMRI results from three male monkeys (Macaca mulatta) during two no-report (only central fixation was required) viewing tasks. One was an abstract sequence viewing task (abbreviated SEQ hereafter) that contained a visual sequence rule and structured timing (as reported in Yusif Rodriguez, et al., 2023). The second task did not contain abstract visual sequences (abbreviated NoSEQ hereafter) but maintained the same stimulus frequencies and periodicity structure as SEQ (**Figure 1**). Our main goals were to 1) test if sequential responses differ between the fundus and more ventral LPFC subregions, and 2) test if and how these subregions respond to changes in the components of abstract visual sequences.

### The fundus of area 46 differentially represents changes to abstract visual sequences

To address our first question, we first tested if sequential responses differed between LPFC subregions: one in the fundus and one adjacent and more ventral. To accomplish this goal, we first needed to define two things: the precise anatomical locations of the regions of interest (ROIs) and sequential responses.

To define the ROIs, we used a parcellation of PFC in conjunction with our previously defined ROI in Yusif Rodriguez (2023). The previous ROI was a 3 mm radius (895 voxels) sphere in the right hemisphere (named R46) based on coordinates chosen for their functional connectivity similarity to human sequence responsive areas of the lateral frontal cortex (Sallet et al., 2013). This ROI was not conducive to comparing directly adjacent areas because of its spherical shape and because it necessarily included some white matter. Therefore, we created ROIs using the MEBRAINS Multilevel Macaque Brain Atlas (Rapan et al., 2023; Balan et al., 2024) that parcellated PFC according to cytoarchitectonics, functional connectivity, and neurochemical data. This atlas divides area 46 into eight distinct regions (four anterior and four posterior) that are then divided into dorsal and ventral shoulder and fundus regions (**Figure 2A**, **B**). Of the area 46 subdivisions, the posterior fundus (p46df and p46vf) regions showed the greatest overlap with the previous R46 ROI constructed from functional connectivity seed coordinates (Sallet et al., 2013; Yusif Rodriguez et al., 2023) (**Figure 2C**). Of the 895 voxels in the previous R46 ROI, 40.5% (420) overlapped with cortical gray matter, and all those voxels overlapped with p46df and p46vf combined. We therefore combined the fundus regions and focused our analyses on the posterior fundus (p46f). We compared p46f (1036 voxels) to an adjacent, more ventral subregion: posterior ventral (p46v, 1060 voxels). We focused on ROIs in the right hemisphere because previous results were observed on the right (though the right hemisphere was not statistically different from the left) (Yusif Rodriguez et al., 2023).

To define sequential responses, we used deviations from an established (habituated) abstract visual sequence in the SEQ task (as in Yusif Rodriguez et al. (2023)). The general logic was that regions that responded to changes in the abstract visual sequence may play a role in tracking that information. To create these changes, monkeys were first habituated to four-item sequences of images that followed a particular rule, e.g., same, same, same, different. Images were drawn from a pool of habituation images. In subsequent blocks, some of the sequences (6 out of 30) were deviant sequences that drew images from a separate deviant pool and differed from habituation sequences in one of four ways: new items, same rule (NISR), rule deviants, number deviants, or double deviants (included to counterbalance the design but not included for analysis). All comparisons to determine abstract sequence responses in the SEQ task were between NISR and rule or number deviants. This comparison controlled for the use of less frequent deviant images and any changes observed would be due to changes in the abstract sequence.

For all analyses, we measured the cerebral blood volume (CBV) of a contrast agent, monocrystalline iron oxide nanoparticle (MION) activity as our indicator of neural activity. We created a single model for both the SEQ and NoSEQ tasks (see Methods for details). Statistical testing was performed on approximately 20-run bins (n = 11), each consisting of data from a single monkey. For each condition, t-values were extracted for that condition compared to baseline (e.g., Rule Deviant > Baseline) to account for potential differences in variance across conditions. These values were used to examine ROI activity throughout, and we refer to comparisons by the condition of interest (i.e., without listing the contrast over baseline, e.g., Rule Deviant). All statistical tests on ROIs were performed on binned data and included a covariate for monkey identity (n = 3). While we report the effect of variation between monkeys in the following analyses, the main focus of the study was not on individual differences, and our discussion focuses on condition effects.

We first tested the hypothesis that sequential responses differ between right p46f and p46v. As in our previous study, we compared the sequence deviants Rule and Number to NISR, with increased activity for deviants indicating sequential processing. Replicating previous results (with the newly defined ROI), both deviant responses were significantly greater than NISR in right p46f (Rule > NISR: p = 0.01; Number > NISR: p = 0.04, **Figure 3A,B**, **Table 3**). In contrast, responses in p46v did not differ between deviants and NISR, resulting in a significant interaction between the two areas for number deviants compared to NISR (p = 0.02, **Figure 3B**, **Table 4**) and a marginal interaction for rule deviants compared to NISR (p = 0.09, **Figure 3A**, **Table 4**). These ROI results were supported by whole brain contrasts of Rule Deviants > NISR (**Figure 3C**) and Number Deviants > NISR (**Figure 3D**) in the SEQ task (**Table 5**). Both deviant types showed significant clusters of activation in right p46f. Therefore, these results support the hypothesis that sequential responses differ between p46f, which responds to changes in abstract visual sequences, and p46v, which does not.

**Figure 3.**
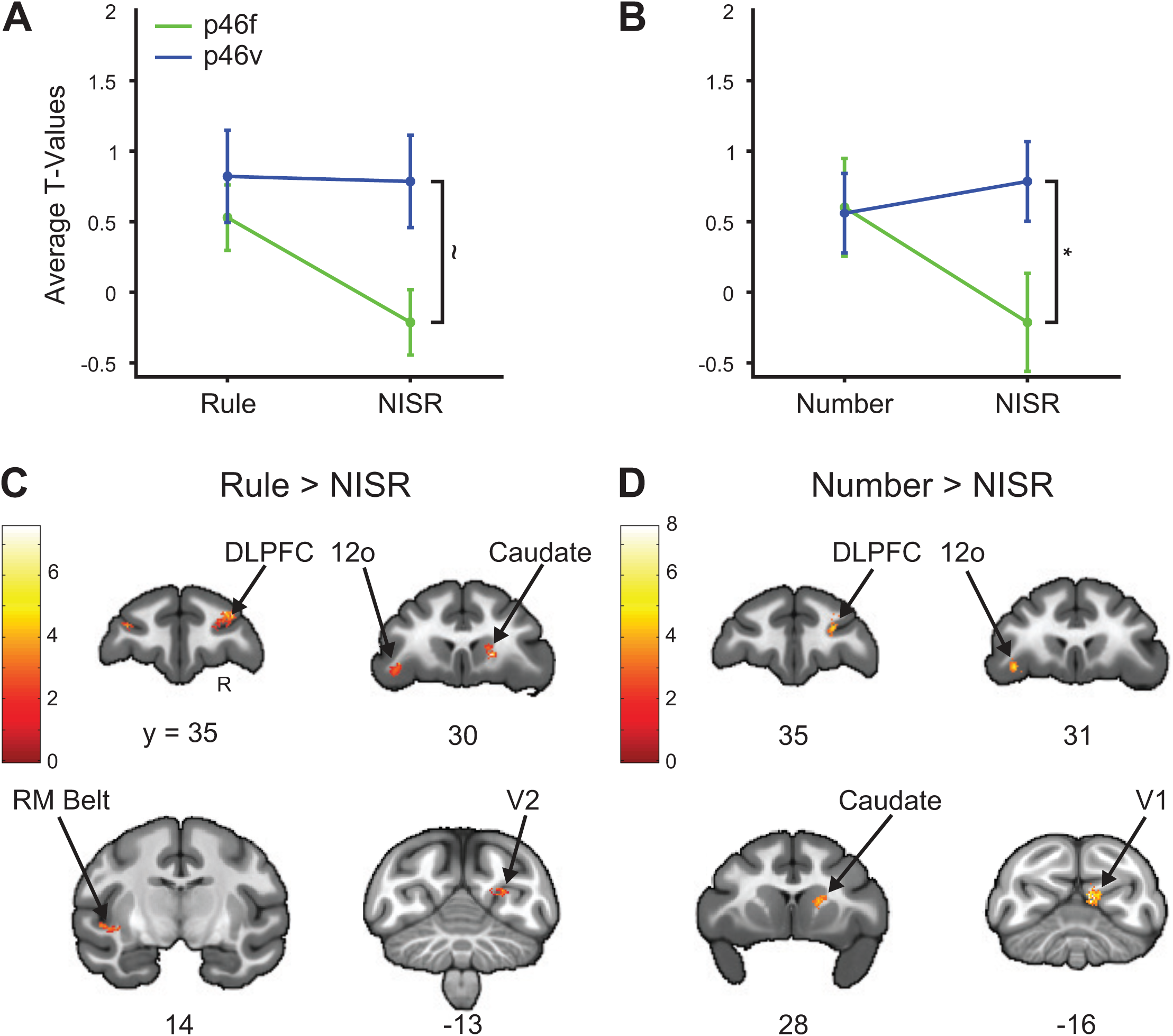
Right p46f and not p46v showed deviant responses in SEQ task. (**A and B)** T-values for the condition of interest > baseline are shown. Error bars are 95% confidence intervals (1.96 x standard error of the within-bin mean). **A**. Responses in rule deviants compared to new items, same rule (NISR) were different between p46f and p46v such that there was a significant main effect of ROI, and a marginal interaction between ROI and condition (indicated with ∼). **B**. Number deviants compared to NISR between p46f and p46v showed a marginal main effect of ROI and significant interaction between ROI and condition. **C**. Voxel-wise contrast of Rule Deviants > NISR, false discovery rate (FDR) error cluster corrected for multiple comparisons (FDRc < 0.05, height p < 0.005 unc., extent = 100) are shown. **D.** Voxel-wise contrast of Number Deviants > NISR (FDRc < 0.05, height p < 0.005 unc., extent = 133) is shown. Color bar indicates T-values in **C** and **D**.

**Table 3.** Activity during SEQ task deviants compared to NISR in right area 46 using repeated measures ANOVAs. P-values in bold are conditions of interest.

| **Rule Deviant** |  |  |  |  |  |
| --- | --- | --- | --- | --- | --- |
| | Factor | DFs | F | p | $\eta_p^2$ |
| 46f | monkey | 2,8 | 1.2 | 0.36 | 0.22 |
|  | condition | 1,8 | 12.3 | **0.01** | 0.61 |
|  | monkey:condition | 2,8 | 4.6 | 0.05 | 0.54 |
| 46v | monkey | 2,8 | 4.5 | 0.05 | 0.53 |
|  | condition | 1,8 | 0 | 0.93 | 0 |
|  | monkey:condition | 2,8 | 0.2 | 0.82 | 0.05 |
| **Number Deviant** |  |  |  |  |  |
| | Factor | DFs | F | p | $\eta_p^2$ |
| 46f | monkey | 2,8 | 0.6 | 0.57 | 0.13 |
|  | condition | 1,8 | 5.8 | **0.04** | 0.42 |
|  | monkey:condition | 2,8 | 5.5 | 0.03 | 0.58 |
| 46v | monkey | 2,8 | 2.8 | 0.12 | 0.41 |
|  | condition | 1,8 | 0.3 | 0.58 | 0.04 |
|  | monkey:condition | 2,8 | 0.1 | 0.88 | 0.03 |

**Table 4.** Comparisons of activity in right p46f and p46v during deviants compared to NISR in the SEQ task using repeated measures ANOVAs. P-values in bold are conditions of interest.

| **Rule Deviant** |  |  |  |  |
| --- | --- | --- | --- | --- |
| Factor | DFs | F | p | $\eta_p^2$ |
| monkey | 2,18 | 0.2 | 0.82 | 0.02 |
| ROI | 1,18 | 5 | 0.04 | 0.22 |
| condition | 1,18 | 2.9 | 0.11 | 0.14 |
| monkey:condition | 2,18 | 1.9 | 0.17 | 0.18 |
| ROI:condition | 1,18 | 3.3 | **0.09** | 0.15 |
| **Number Deviant** |  |  |  |  |
| Factor | DFs | F | p | $\eta_p^2$ |
| monkey | 2,18 | 0.6 | 0.57 | 0.06 |
| ROI | 1,18 | 3.7 | 0.07 | 0.17 |
| condition | 1,18 | 1 | 0.32 | 0.05 |
| monkey:condition | 2,18 | 2.5 | 0.11 | 0.22 |
| ROI:condition | 1,18 | 6 | **0.02** | 0.25 |

**Table 5.** Coordinates of activity clusters in Rule and Number deviant > NISR contrasts in the SEQ task.

| Contrast Location | Extent<br>(vox) | Peak T-val | X | Y | Z |
| --- | --- | --- | --- | --- | --- |
| **Rule Deviant > NISR** |  |  |  |  |  |
| Rostral Medial Frontal Pole | 105 | 6.25 | 0.5 | 45.5 | 14.5 |
| Dorsal Area 46 | 381 | 7.53 | 14 | 35.5 | 25.5 |
| Ventral Area 46 | 100 | 5.53 | -15 | 35.5 | 22 |
| Medial Agranular Insular Region | 159 | 7.53 | 8 | 30.5 | 15 |
| Orbital Area 12 | 148 | 5.59 | -18 | 30.5 | 12.5 |
| Area F5 of Ventral Pre-motor Cortex | 173 | 6.06 | -19 | 13 | 9.5 |
| Granular Layer of Dentate Gyrus | 137 | 5.66 | -9.5 | 2.5 | 12.5 |
| Cerebellum | 229 | 4.85 | 6 | -5.5 | 5.5 |
| Visual Area 2 | 123 | 5.65 | 12 | -12 | 20 |
|  | 115 | 7.21 | 3 | -22 | 23 |
| **Number Deviant > NISR** |  |  |  |  |  |
| Dorsal Area 46 | 166 | 5.73 | 11 | 36.5 | 25.5 |
| Orbital Area 12 | 133 | 5.17 | -18 | 31 | 10.5 |
| Medial Area 13 | 162 | 5.21 | 7.5 | 27.5 | 17.5 |
| Visual Area 2 | 332 | 7.99 | 5 | -17 | 18 |

### The ventral shoulder of area 46 differentially represents changes to abstract sequence components

We next addressed the second question of if and how these regions respond to changes in sequential components in the absence of an abstract visual sequence. We examined two main features in the NoSEQ task: image periodicity (timing template) and identity. These two image features are components of the SEQ task, exist in parallel in the NoSEQ task, and will be described further below.

To test for responses in the DLPFC related to image periodicity, we compared standard to nonstandard timing templates in the NoSEQ task. Previous studies showed that regions of the frontal cortex process different types of timing structures (Onoe et al., 2001; Genovesio et al., 2006; Chiba et al., 2021), raising the possibility that a difference in the timing structure alone could be a component of sequence responses. In the context of this experiment, image periodicity refers to the timing template used. In the SEQ and NoSEQ tasks, most sequences/groups had one of six possible standard timing templates (referred to as habituation in the SEQ task, **Figure 1**).

A unique timing template (0.2 s image duration for medium, 1.7 s, total duration) was used infrequently for deviants in the SEQ task (after the first block, 6 out of 30 sequences). In the NoSEQ task that structure was mirrored: after the first block, 6 out of the 30 stimulus groupings used a nonstandard timing template. These blocks either contained six 4-image nonstandard timing (4NST) or three each of 2– and 6-image nonstandard timing (2/6NST). Importantly, even though the timing template was the same as for SEQ deviants (just termed differently for the NoSEQ task), the images in NoSEQ were pseudorandomly presented and were not composed of entirely nonstandard images. To determine if brain areas responded to changes in timing template alone, we compared responses to these nonstandard timing templates to the six other standard timing templates in the NoSEQ task.

We tested the hypothesis that p46v would show a greater difference in responses to image periodicity than p46f in the NoSEQ task. In general, more ventral LPFC regions are thought to have more object-based or visual responses (Meyer et al., 2011; Yamagata et al., 2012; Tang et al., 2021; Xu et al., 2022). First, we found that changes in timing structure alone did not elicit deviant responses in right p46f. There were no reliable differences between 4NST (p = 0.86, **Figure 4A**, **Table 6**) or 2/6NST compared to standard timing templates (p = 0.85, **Figure 4B**, **Table 6**). In contrast, right p46v showed reliable differences when comparing standard to 4NST (p = 0.03, **Figure 4A**, **Table 6**) and standard to 2/6NST (p = 0.01, **Figure 4B**, **Table 6**). When directly comparing p46f and p46v, overall responses were greater in p46v than p46f (p = 0.01, **Table 7**) and there was a significant interaction such that the difference between nonstandard and standard was significantly different by ROI for 2/6NST and marginal for 4NST (ROI x condition, 2/6NST: p = 0.05, 4NST: p = 0.09; **Table 7**). These results were supported by whole brain contrasts of 4NST > Standard Timing and 2/6NST > Standard Timing showing no significant clusters of activation in p46f with other significant activation in distinct visual and association areas (**Figure 4C,D**, **Table 8**). Together, these results support the hypothesis that p46v responds to image periodicity and dissociates from responses in p46f. Further, p46v may be part of a network of other brain areas that is specialized to detect periodicity differences, independent of abstract sequential structure.

**Figure 4.**
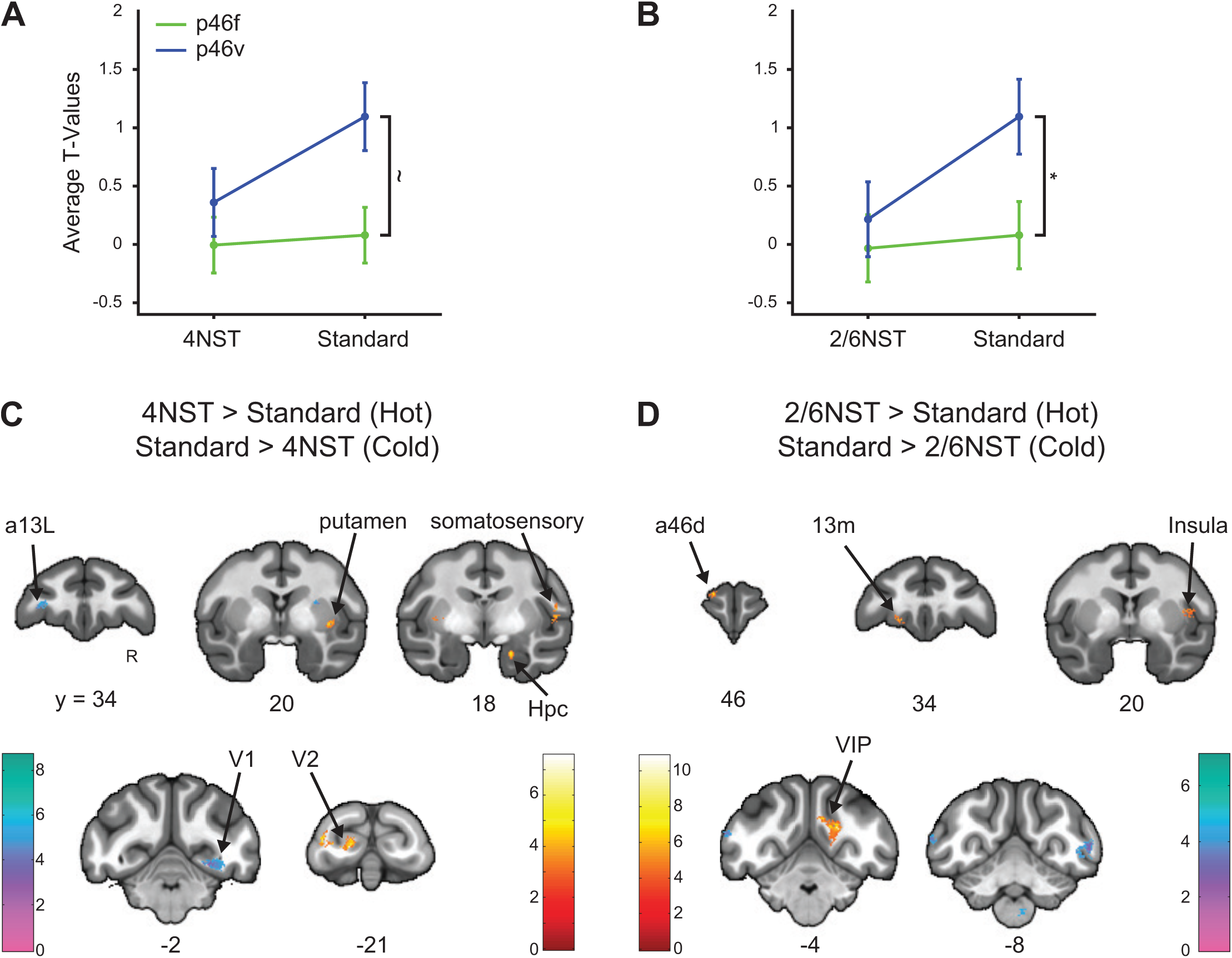
Right p46v and not p46f showed different responses to timing templates in NoSEQ. (**A and B)** T-values for the condition of interest > baseline are shown. Error bars are 95% confidence intervals (1.96 x standard error of the within-bin mean). **A.** 4NST compared to standard timing between p46f and p46v shows a significant main effect of ROI and a marginal interaction between ROI and condition (indicated with ∼). **B**. 2/6NST compared to standard between p46f and p46v shows a significant main effect of ROI and interaction between ROI and condition (indicated with *). **C.** Voxel wise contrasts of 4NST > Standard timing (Hot colors) false discovery rate (FDR) error cluster corrected for multiple comparisons (FDRc < 0.05, height p < 0.005 unc., extent = 84), overlaid with Standard timing > 4NST (Cold colors) false discovery rate (FDR) error cluster corrected for multiple comparisons (FDRc < 0.05, height p < 0.005 unc., extent = 204) are shown. **D.** Voxel wise contrasts of 2/6NST > Standard timing (hot colors; FDRc < 0.05, height p < 0.005 unc., extent = 94), overlaid with Standard timing > 2/6NST (cold colors; FDRc < 0.05, height p < 0.005 unc., extent = 104) are shown. Color bars indicate T-values in **C** and **D**.

**Table 6.** Activity during NoSEQ task 4NST and 2/6NST compared to standard timing in right area 46 using repeated measures ANOVAs. P-values in bold are conditions of interest.

| **4NST** |  |  |  |  |  |
| --- | --- | --- | --- | --- | --- |
|  | Factor | DFs | F | p | eta2p |
| 46f | monkey | 2,8 | 0.8 | 0.5 | 0.16 |
|  | condition | 1,8 | 0 | 0.86 | 0 |
|  | monkey:condition | 2,8 | 0.1 | 0.95 | 0.01 |
| 46v | monkey | 2,8 | 9.6 | 0.01 | 0.71 |
|  | condition | 1,8 | 6.8 | **0.03** | 0.46 |
|  | monkey:condition | 2,8 | 5.1 | 0.04 | 0.56 |
| **2/6NST** |  |  |  |  |  |
|  | Factor | DFs | F | p | eta2p |
| 46f | monkey | 2,8 | 0.1 | 0.95 | 0.01 |
|  | condition | 1,8 | 0 | 0.85 | 0 |
|  | monkey:condition | 2,8 | 1.1 | 0.37 | 0.22 |
| 46v | monkey | 2,8 | 14.5 | 0 | 0.78 |
|  | condition | 1,8 | 10 | **0.01** | 0.55 |
|  | monkey:condition | 2,8 | 4.4 | 0.05 | 0.52 |

**Table 7.** Comparisons of activity in right p46f and p46v during nonstandard compared to standard timing in the NoSEQ task using repeated measures ANOVAs. P-values in bold are conditions of interest.

| **4NST** |  |  |  |  |
| --- | --- | --- | --- | --- |
| Factor | DFs | F | p | $\eta_p^2$ |
| monkey | 2,18 | 6.3 | 0.01 | 0.41 |
| ROI | 1,18 | 8.4 | **0.01** | 0.32 |
| condition | 1,18 | 2.8 | 0.11 | 0.14 |
| monkey:condition | 2,18 | 1.9 | 0.19 | 0.17 |
| ROI:condition | 1,18 | 3.1 | **0.09** | 0.15 |
| **2/6NST** |  |  |  |  |
| Factor | DFs | F | p | $\eta_p^2$ |
| monkey | 2,18 | 3.4 | 0.06 | 0.27 |
| ROI | 1,18 | 8.2 | **0.01** | 0.31 |
| condition | 1,18 | 5.3 | 0.03 | 0.23 |
| monkey:condition | 2,18 | 5 | 0.02 | 0.36 |
| ROI:condition | 1,18 | 4.3 | **0.05** | 0.19 |

**Table 8.**
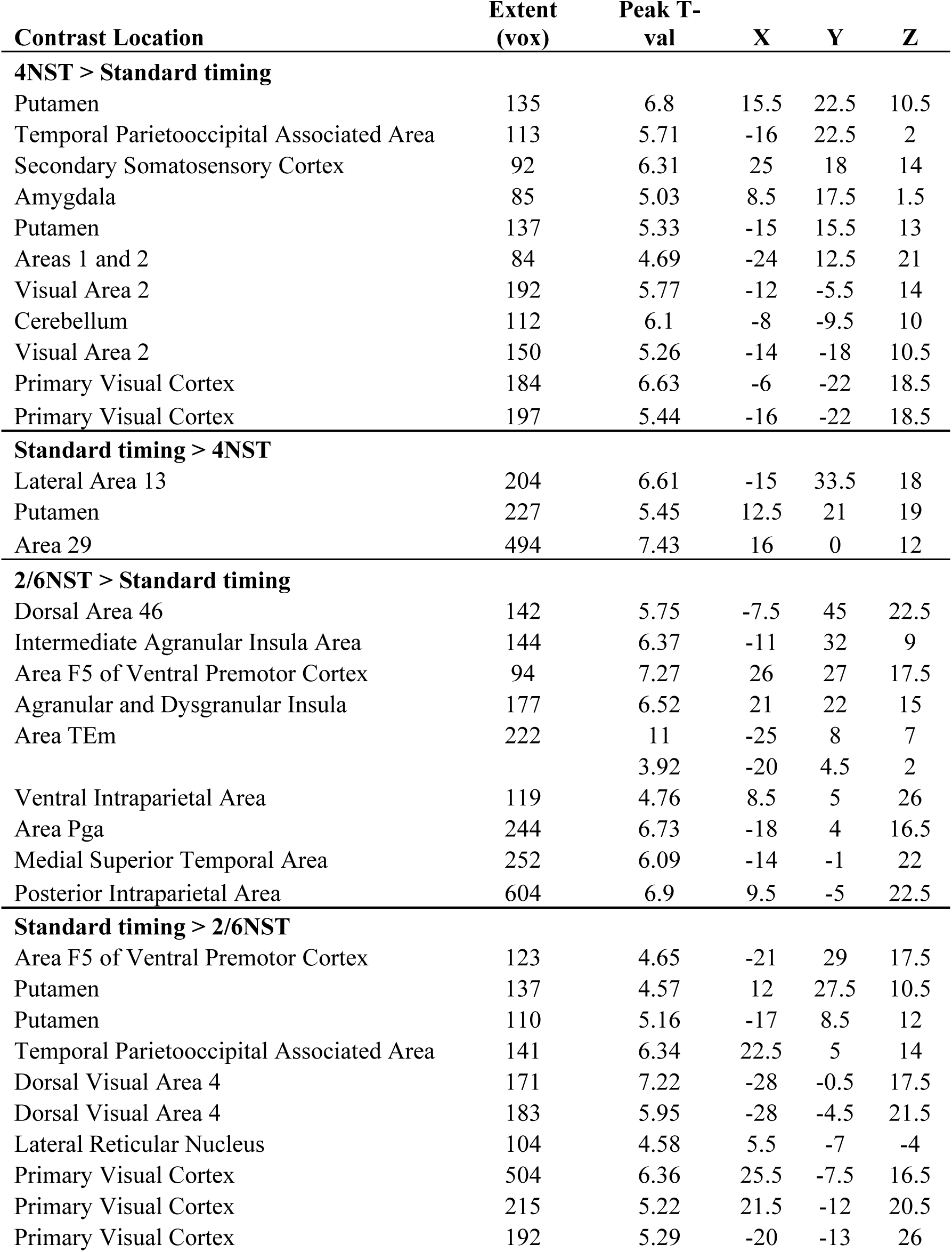
Coordinates of activity clusters in 4NST and 2/6NST > Standard timing contrasts in the NoSEQ task.

To test for responses in the LPFC related to image identity, we compared standard to nonstandard images in the NoSEQ task. Deviant/nonstandard images in the SEQ task were less frequent. It is unlikely that the deviant responses observed in the SEQ task were driven by these images, because deviant comparisons were all made across conditions that contained images from the deviant pool (e.g., rule deviant vs. NISR, **Figure 1**). However, infrequent or surprising images have been shown to drive responses in LPFC (Chao et al., 2018; Camalier et al., 2019; Grohn et al., 2020). Therefore, we aimed to determine if responses in area 46 could be driven by less frequent image presentations, independent of sequential context. To examine this sequence component, we again used conditions that were separate from an abstract visual sequence, i.e., in the NoSEQ task. We compared responses to the randomly interspersed nonstandard images to standard images to ensure other aspects of the task were held constant.

We tested the hypothesis that p46v would show a greater difference in responses to image identity than p46f in the NoSEQ task. First, we found that nonstandard responses were not reliably different from standard image responses in right p46f (p = 0.67, **Figure 5A**, **Table 9**). In contrast, p46v differentiated between standard and nonstandard images, but with reliably greater responses for standard images (p = 0.01, **Figure 5A**, **Table 9**). We directly compared responses in p46f to p46v and found that responses to nonstandard compared to standard images were reliably different (ROI x condition: p = 0.03, **Table 10**). These ROI results showing greater responses to standard than nonstandard images were also supported by whole brain contrasts (**Figure 5B**). Significant clusters of activation were observed for Standard > Nonstandard images in right p46v but no other area 46 subregions (**Table 11**). In Nonstandard > Standard image there were no significant clusters in the frontal cortex. These results support the hypothesis that p46v differentially represents standard and nonstandard images, and p46f does not.

**Figure 5.**
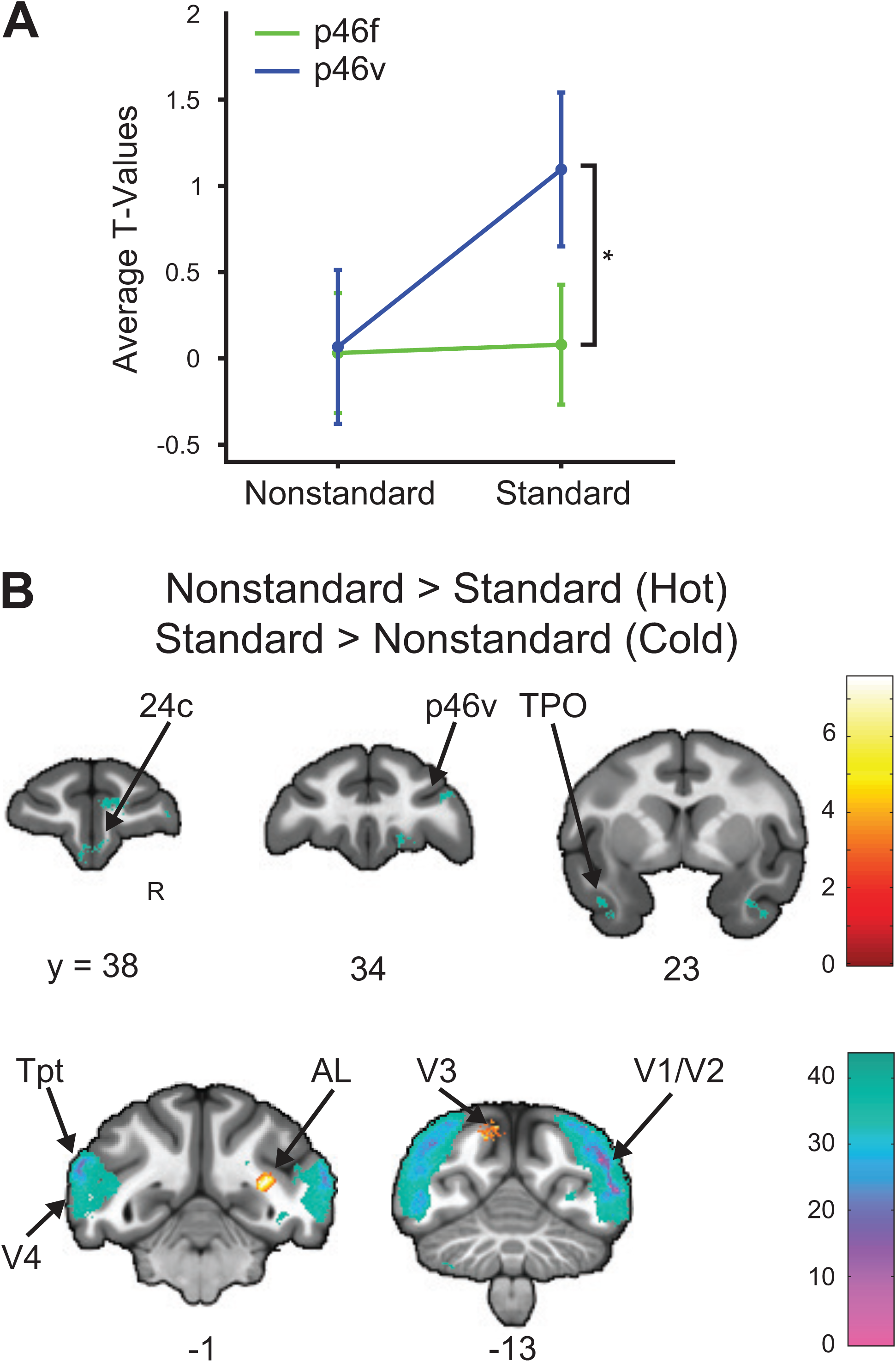
Right p46v and not p46f shows different responses to standard and nonstandard images in the NoSEQ task. T-values for the condition of interest > baseline are shown. Error bars are 95% confidence intervals (1.96 x standard error of the within-bin mean). **A.** Nonstandard compared to standard images showing reliable differences between p46f and p46v showing a significant main effect of ROI with a significant interaction of ROI and condition (indicated with *). **B.** Voxel wise contrasts of Nonstandard > Standard images false discovery rate (FDR) error cluster corrected for multiple comparisons (hot colors, FDRc < 0.05, height p < 0.005 unc., extent = 101), overlaid with Standard > Nonstandard image (cold colors, FDRc < 0.05, height p < 0.005 unc., extent = 88) are shown. Color bar indicates T-values.

**Table 9.** Activity during NoSEQ task nonstandard images compared to standard images in right area 46 using repeated measures ANOVAs. P-values in bold are conditions of interest.

| | Factor | DFs | F | p | $\eta_p^2$ |
| --- | --- | --- | --- | --- | --- |
| 46f | monkey | 2,8 | 1.3 | 0.32 | 0.25 |
|  | condition | 1,8 | 0.2 | 0.67 | 0.02 |
|  | monkey:condition | 2,8 | 3.3 | 0.09 | 0.45 |
| 46v | monkey | 2,8 | 9.7 | 0.01 | 0.71 |
|  | condition | 1,8 | 11.8 | **0.01** | 0.6 |
|  | monkey:condition | 2,8 | 10.2 | 0.01 | 0.72 |

**Table 10.** Comparisons of activity in right p46f and p46v during standard compared to nonstandard images in the NoSEQ task using repeated measures ANOVAs. P-values in bold are conditions of interest.

| Factor | DFs | F | p | $\eta_p^2$ |
| --- | --- | --- | --- | --- |
| monkey | 2,18 | 1.8 | 0.2 | 0.17 |
| ROI | 1,18 | 8.7 | 0.01 | 0.33 |
| condition | 1,18 | 7 | 0.02 | 0.28 |
| monkey:condition | 2,18 | 12 | 0 | 0.57 |
| ROI:condition | 1,18 | 6 | **0.03** | 0.25 |

**Table 11.**
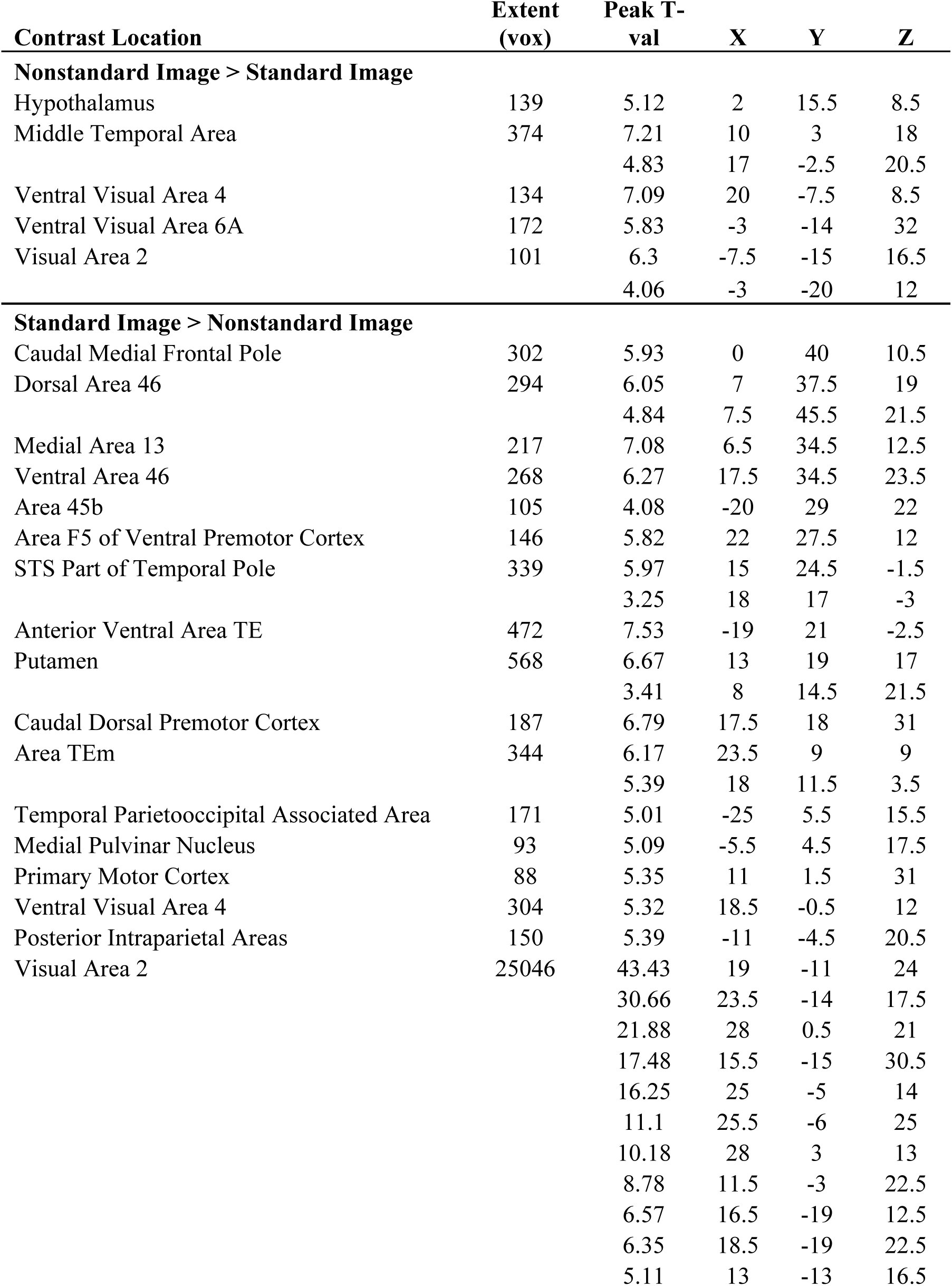
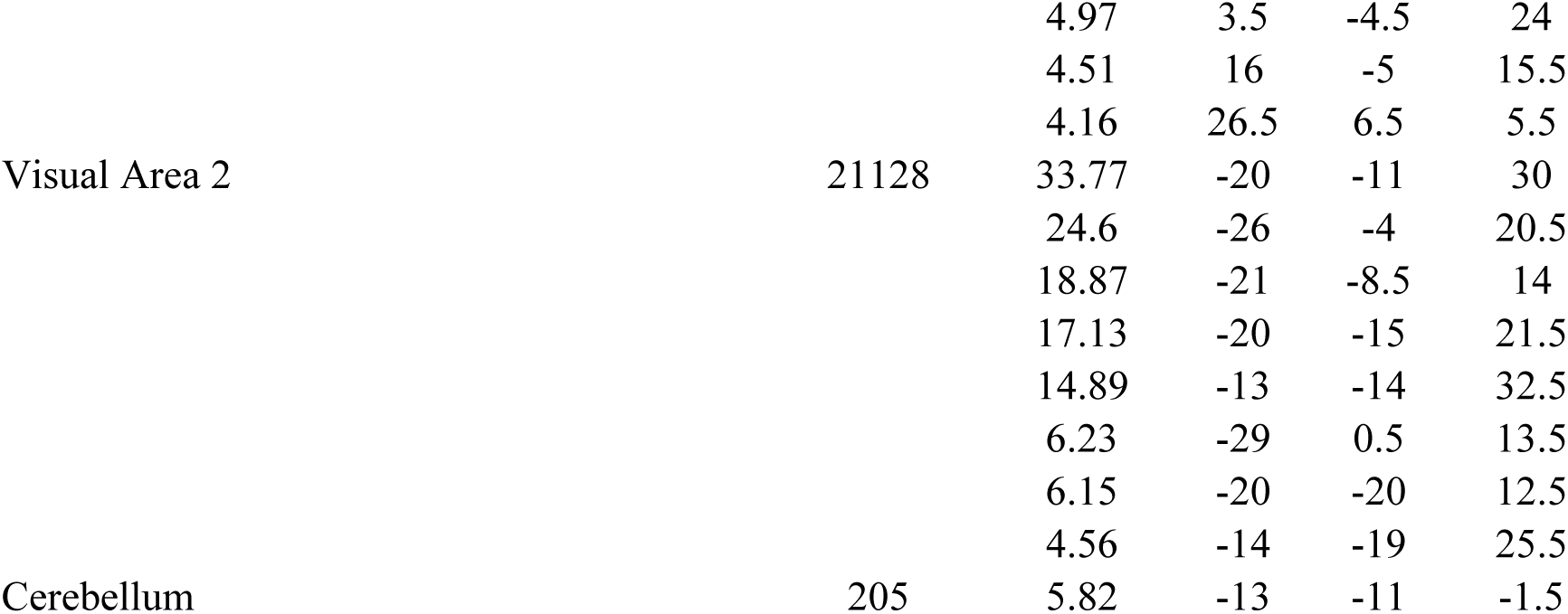
Coordinates of activity clusters in Nonstandard Image > Standard Image and Standard Image > Nonstandard Image contrasts in the NoSEQ task.

## Discussion

We had two main goals in this experiment: 1) to test if SEQ deviant responses previously observed in p46f were different from those in p46v, and 2) to test if and how p46f and p46v responded to changes in abstract visual sequence components, image identity and periodicity, alone in a NoSEQ task. We hypothesized that p46v would not respond to changes in abstract visual sequences as a whole, but to the defined components. The results generally supported the hypotheses, with p46f responding more strongly than p46v to abstract sequence deviants in the SEQ task but not to differences in sequential components (standard vs nonstandard) in NoSEQ, further strengthening its role as an area that uniquely represents abstract sequential changes. P46v instead differentiated between standard and nonstandard sequence components, image identity and periodicity, in NoSEQ. This result supported our predictions that this region was more influenced by visual sensory inputs and not necessarily the higher order structures in abstract sequences. These results provide important knowledge of the functional subdivisions within area 46 of the LPFC and scaffold future understanding of this important area for cognition.

The observed differences between p46f and p46v may expand on the notion of “dorsal” and “ventral” distinctions within the LPFC. Classic anatomical definitions of dorsal and ventral have used the principal sulcus as a dividing line, bisecting the fundus between the two (e.g., Petrides and Pandya (1999)). Extracellular electrophysiology experiments often focus on the cortical surface of LPFC, understandably due to the ability to visualize locations and position electrodes and arrays. Therefore, it is less clear how the fundus region itself may or may not fit into functional generalizations of LPFC. Anatomy and the present experiment suggest a more distinct functional role for p46f, at least within more posterior LPFC regions. The present results strikingly align with a multimodal parcellation of macaque area 46 showing that p46f hierarchically clustered with the most rostral regions of LPFC whereas p46v clustered with more caudal sensory and motor areas (Rapan et al., 2023). Current schema of the functional organization of LPFC suggest that more spatial and action-oriented responses are localized more dorsally and more non-spatial and object-oriented responses are localized more ventrally (Meyer et al., 2011; Yamagata et al., 2012; Tang et al., 2021; Xu et al., 2022). These results are also broadly consistent with a proposed distinction of dorsal ‘How’ (perception to action transformation) and ventral ‘What’ (identity) information in the LPFC (O’Reilly, 2010), if it is considered as a gradient and “actions” could be considered non-motor. Further, we did not observe responses to changes in abstract visual sequences in p46v, as others have observed in what was termed VLPFC (Wang et al., 2015; Bellet et al., 2024). It is unclear if p46v and VLPFC are anatomically overlapping. More experiments are needed before strong conclusions can be drawn, but the results here suggest that the subregions within what has been referred to as DLPFC and VLPFC should be carefully considered, and the fundus may need separate classification. Future experiments could benefit from recent developments in whole brain imaging technology including functional contrast agents, PET-MR, and high field fMRI to localize anatomical and functional brain regions both independently and pre/post-electrophysiological recordings.

We did not observe responses in p46f to changes in the periodicity alone in NoSEQ. In contrast, right p46v showed differences in responding to standard and nonstandard timing templates, with increased responses to the more frequent standard timing presentations. These observations are generally consistent with previous observations of timing-related activity in monkey LPFC (Niki and Watanabe, 1979; Onoe et al., 2001; Genovesio et al., 2006; Cueva et al., 2020; Chiba et al., 2021), although the precise anatomical location was not specified. Human LPFC responses in temporal expectation tasks (Coull and Nobre, 2008) were also similar. Outside of area 46 we observed some of the same regions that have been observed for duration perception in monkeys, such as putamen, cerebellum, and V2 (Onoe et al., 2001). We also observed regions similar to those observed in humans related to temporal expectation such as the basal ganglia, temporal cortex, and cerebellum (Coull and Nobre, 2008). Together these results illustrate the specificity of subregions within area 46 and suggest that adjacent subregions code for different stimulus properties.

Area p46v responses differentiated between standard and nonstandard images in NoSEQ while p46f did not. Responses in p46v were greater to standard images than nonstandard images. These results suggest that this response was not a typical ‘surprise’ response. In our task, standard images were presented with greater frequency, and closer in time to each other, than the isolated individual appearances of the nonstandard images. Therefore, such responses could be partially driven by the increased frequency, and thus a greater summed BOLD response, of standard compared to nonstandard images and. This increased image frequency may have been the primary driver of responses in regions such as visual cortex, which showed relatively large significant clusters in Standard > Nonstandard NoSEQ images. However, if frequency was the only driver of such responses, then we would have likely observed these responses throughout the whole brain rather than in a specific set of regions. Outside of the visual cortex there were roughly equal numbers of areas that showed responses to the reverse contrast (Nonstandard Image > Standard Image), but these responses were not located in area 46. An intriguing possibility is that a greater response to the standard images in p46v is due to the previous association that those images have to abstract visual sequences, or their greater familiarity (Rainer et al., 1999; Stern et al., 2001; Leaver et al., 2009). Further investigation will be necessary to determine if that association is a component of the response in p46v along with regions observed for Standard Image > Nonstandard Image in the whole brain. There were a small number of regions observed outside of the frontal cortex that showed significantly greater responses to nonstandard images. These regions were not necessarily overlapping with areas typically associated with ‘surprise’ or prediction error (Grohn et al., 2020), again raising the prospect that a form of association may govern these responses as well. Further research will be needed to discern the underlying driving forces, but the fact remains that sensory related responses localize to p46v and not adjacent p46f, again illustrating the specificity of responses within area 46.

This study’s approach was limited in the following ways. First, though p46f and p46v showed significant differences, some effect sizes remained small. These results could be related to limitations of whole-brain event-related monkey fMRI: the spatial resolution, signal-to-noise, and inherent smoothness of the data. There could be differences in alignment of the voxels with the regions and partial volume effects that would be difficult to resolve without fundamentally changing the experiment by scanning a small volume at higher resolution, using a greater field strength (which may not be available), or greatly increasing the sample size (introducing other limitations). However, these experiments provide an ideal foundation for techniques with higher spatial resolution such as electrophysiological recordings. Second, while the no-report paradigm confers the advantage of eliminating possible confounds due to executing responses, it does not allow for direct comparisons with behavior performance. In other words, even though we have observed that regions within area 46 respond to changes in stimuli, we do not know how such information may contribute to decisions or the production of actions. Third, we have specifically focused on two subregions within area 46, which is itself only one of several areas defined as belonging to LPFC. Other regions within and beyond LPFC warrant further investigation, and the results here potentially contextualize further differences within and among subregions. Though some of the preceding items are limitations of the chosen task and technique, we hope that this experiment and others like it highlights the utility of different data acquisition modalities and opens important avenues of future research.

In conclusion, we provide unique evidence for the anatomical and functional specificity of abstract visual sequence deviant responses in a specific subregion of LPFC, p46f. In tandem, we provide evidence that the adjacent region p46v, differentiates image identity and periodicity components. These results reinforce the potential parallel with findings in human brain areas. Rostrolateral PFC in humans is necessary for abstract task sequences and is most analogous to p46f in monkeys (Desrochers et al., 2015; Yusif Rodriguez et al., 2023). Further, these results illustrate the utility in using fMRI to isolate components to cognitive processes, in this case, sequential components. The LPFC, which may have in the past appeared to be a more homogenous region, may in fact be even more distinct in its subdivisions and functional mapping. This study lays the foundation for an approach to functionally dissociating subregions in the cortical structures that underlie many complex and abstract daily functions, such as cooking a meal or appreciating a piece of music.

## Acknowledgements

This study was supported by the National Science Foundation (NSF) Established Program to Stimulate Competitive Research (EPSCoR) Neural Basis of Attention Grant 1632738 (N.Y.R. and T.M.D.), the National Institute of General Medical Sciences (NIGMS)-National Institutes of Health (NIH) Initiative to Maximize Student Development Grant IMSD R25GM083270 (N.Y.R.), the NIH-NIGMS Grant COBRE P20GM103645 (T.M.D), NIH National Institutes of Mental Health (NIMH) R21MH125010 (T.M.D.), NSF Faculty Early Career Development (CAREER) Program Award BCS-2143656 (T.M.D.), NIMH Research Project Grant R01MH131615 (T.M.D.), and the Carney Institute for Brain Science Innovation Award (T.M.D.). Part of this research was conducted using computational resources and services at the Center for Computation and Visualization, Brown University (NIH Grant S10OD025181).

We thank Matthew Maestri for his assistance with animal training and data collection. We thank Dr. Michael Worden, Lynn Fanella, Fabienne McEleney, and Brown University’s MRI Facilities staff for their support and guidance throughout this project. We thank Dr. Lucija Jankovic-Rapan, Dr. Nicola Palomero-Gallagher and Dr. Seán Froudist-Walsh for their assistance and sharing of the MEBRAINS atlas and regions of interest used for analysis. We also thank Dr. David Sheinberg, Dr. Amitai Shenhav, Dr. Katherine Conen, and Hannah Doyle for their continued support throughout this project and preparation of this publication as well as members of both the Sheinberg and the Desrochers Labs for many helpful discussions and contributions.

## Author Contributions

Nadira Yusif Rodriguez: Conceptualization, Investigation, Formal analysis, Writing – Original Draft, Writing – Review & Editing, Visualization. Aarit Ahuja: Investigation, Writing – Review & Editing. Debaleena Basu: Conceptualization, Investigation, Writing – Review & Editing. Theresa H. McKim: Conceptualization, Investigation, Writing – Review & Editing. Theresa M. Desrochers: Conceptualization, Methodology, Investigation, Writing – Original Draft, Writing – Review & Editing, Supervision, Resources, Funding acquisition.

## References

1. Balan PF, Zhu Q, Li X, Niu M, Rapan L, Funck T, Wang H, Bakker R, Palomero-Gallagher N, Vanduffel W (2024) MEBRAINS 1.0: A new population-based macaque atlas. Imaging Neuroscience 2:1–26.

2. Bellet ME, Gay M, Bellet J, Jarraya B, Dehaene S, van Kerkoerle T, Panagiotaropoulos TI (2024) Spontaneously emerging internal models of visual sequences combine abstract and event-specific information in the prefrontal cortex. Cell Rep 43:113952.

3. Camalier CR, Scarim K, Mishkin M, Averbeck BB (2019) A Comparison of Auditory Oddball Responses in Dorsolateral Prefrontal Cortex, Basolateral Amygdala, and Auditory Cortex of Macaque. Journal of Cognitive Neuroscience 31:1054–1064.

4. Chao ZC, Takaura K, Wang L, Fujii N, Dehaene S (2018) Large-Scale Cortical Networks for Hierarchical Prediction and Prediction Error in the Primate Brain. Neuron 100:1252–1266.e3.

5. Chiba A, Morita K, Oshio K, Inase M (2021) Neuronal activity in the monkey prefrontal cortex during a duration discrimination task with visual and auditory cues. Sci Rep 11:17520.

6. Coull JT, Nobre AC (2008) Dissociating explicit timing from temporal expectation with fMRI. Current Opinion in Neurobiology 18:137–144.

7. Cueva CJ, Saez A, Marcos E, Genovesio A, Jazayeri M, Romo R, Salzman CD, Shadlen MN, Fusi S (2020) Low-dimensional dynamics for working memory and time encoding. Proc Natl Acad Sci USA.

8. Desrochers TM, Chatham CH, Badre D (2015) The necessity of rostrolateral prefrontal cortex for higher-level sequential behavior. Neuron 87:1357–1368.

9. Esmailpour H, Raman R, Vogels R (2023) Inferior temporal cortex leads prefrontal cortex in response to a violation of a learned sequence. Cereb Cortex 33:3124–3141.

10. Genovesio A, Tsujimoto S, Wise SP (2006) Neuronal Activity Related to Elapsed Time in Prefrontal Cortex. Journal of Neurophysiology 95:3281–3285.

11. Grohn J, Schüffelgen U, Neubert F-X, Bongioanni A, Verhagen L, Sallet J, Kolling N, Rushworth MFS (2020) Multiple systems in macaques for tracking prediction errors and other types of surprise. PLoS Biol 18:e3000899.

12. Jean-Baptiste Poline MB (2002) Region of interest analysis using an SPM toolbox.

13. Kim HF, Hikosaka O (2013) Distinct basal ganglia circuits controlling behaviors guided by flexible and stable values. Neuron 79:1001–1010.

14. Leaver AM, Lare JV, Zielinski B, Halpern AR, Rauschecker JP (2009) Brain Activation during Anticipation of Sound Sequences. J Neurosci 29:2477–2485.

15. Leite FP, Tsao D, Vanduffel W, Fize D, Sasaki Y, Wald LL, Dale AM, Kwong KK, Orban G a, Rosen BR, Tootell RBH, Mandeville JB (2002) Repeated fMRI using iron oxide contrast agent in awake, behaving macaques at 3 Tesla. NeuroImage 16:283–294.

16. McLaren DG, Kosmatka KJ, Oakes TR, Kroenke CD, Kohama SG, Matochik JA, Ingram DK, Johnson SC (2009) A population-average MRI-based atlas collection of the rhesus macaque. NeuroImage 45:52–59.

17. Meyer T, Qi X-L, Stanford TR, Constantinidis C (2011) Stimulus Selectivity in Dorsal and Ventral Prefrontal Cortex after Training in Working Memory Tasks. J Neurosci 31:6266– 6276.

18. Miyashita Y, Higuchi SI, Sakai K, Masui N (1991) Generation of fractal patterns for probing the visual memory. Neuroscience Research 12:307–311.

19. Niki H, Watanabe M (1979) Prefrontal and cingulate unit activity during timing behavior in the monkey. Brain Res 171:213–224.

20. Onoe H, Komori M, Onoe K, Takechi H, Tsukada H, Watanabe Y (2001) Cortical Networks Recruited for Time Perception: A Monkey Positron Emission Tomography (PET) Study. NeuroImage 13:37–45.

21. O’Reilly RC (2010) The What and How of prefrontal cortical organization. Trends in Neurosciences 33:355–361.

22. Petrides M, Pandya DN (1999) Dorsolateral prefrontal cortex: comparative cytoarchitectonic analysis in the human and the macaque brain and corticocortical connection patterns. Eur J Neurosci 11:1011–1036.

23. Poline J-B, Brett M (2012) The general linear model and fMRI: Does love last forever? NeuroImage 62:871–880.

24. Rainer G, Rao SC, Miller EK (1999) Prospective Coding for Objects in Primate Prefrontal Cortex. J Neurosci 19:5493–5505.

25. Rapan L, Froudist-Walsh S, Niu M, Xu T, Zhao L, Funck T, Wang X-J, Amunts K, Palomero-Gallagher N (2023) Cytoarchitectonic, receptor distribution and functional connectivity analyses of the macaque frontal lobe Badre D, Baker CI, Thiebaut de Schotten M, eds. eLife 12:e82850.

26. Sallet J, Mars RB, Noonan MP, Neubert F-X, Jbabdi S, O’Reilly JX, Filippini N, Thomas AG, Rushworth MF (2013) The organization of dorsal frontal cortex in humans and macaques. J Neurosci 33:12255–12274.

27. Sirmpilatze N, Klink PC (2020) RheMAP: Non-linear warps between common rhesus macaque brain templates. Available at: https://zenodo.org/records/3668510 [Accessed July 10, 2024].

28. Stern CE, Sherman SJ, Kirchhoff BA, Hasselmo ME (2001) Medial temporal and prefrontal contributions to working memory tasks with novel and familiar stimuli. Hippocampus 11:337–346.

29. Suda Y, Tada M, Matsuo T, Kawasaki K, Saigusa T, Ishida M, Mitsui T, Kumano H, Kirihara K, Suzuki T, Matsumoto K, Hasegawa I, Kasai K, Uka T (2022) Prediction-Related Frontal-Temporal Network for Omission Mismatch Activity in the Macaque Monkey. Front Psychiatry 13:557954.

30. Tang H, Bartolo R, Averbeck BB (2021) Reward-related choices determine information timing and flow across macaque lateral prefrontal cortex. Nat Commun 12:894.

31. Uhrig L, Dehaene S, Jarraya B (2014) A Hierarchy of Responses to Auditory Regularities in the Macaque Brain. J Neurosci 34:1127–1132.

32. Vanduffel W, Farivar R (2014) Functional MRI of Awake Behaving Macaques Using Standard Equipment. In: Advanced Brain Neuroimaging Topics in Health and Disease (Papageorgiou TD, Christopoulos GI, Smirnakis SM, eds), pp Ch. 6. Rijeka: IntechOpen. Available at: 10.5772/58281 [Accessed March 9, 2023].

33. Vergnieux V, Vogels R (2020) Statistical Learning Signals for Complex Visual Images in Macaque Early Visual Cortex. Front Neurosci 14:789.

34. Walker AE (1940) A cytoarchitectural study of the prefrontal area of the macaque monkey. Journal of Comparative Neurology 73:59–86.

35. Wang L, Uhrig L, Jarraya B, Dehaene S (2015) Representation of numerical and sequential patterns in macaque and human brains. Current Biology 25:1966–1974.

36. Xu R, Bichot NP, Takahashi A, Desimone R (2022) The cortical connectome of primate lateral prefrontal cortex. Neuron 110:312–327.e7.

37. Yamagata T, Nakayama Y, Tanji J, Hoshi E (2012) Distinct information representation and processing for goal-directed behavior in the dorsolateral and ventrolateral prefrontal cortex and the dorsal premotor cortex. J Neurosci 32:12934–12949.

38. Yusif Rodriguez N, McKim TH, Basu D, Ahuja A, Desrochers TM (2023) Monkey Dorsolateral Prefrontal Cortex Represents Abstract Visual Sequences during a No-Report Task. J Neurosci 43:2741–2755.

